# Prediction of infectivity of SARS-CoV2: Mathematical model with analysis of docking simulation for spike proteins and angiotensin-converting enzyme 2

**DOI:** 10.1101/2022.03.30.486373

**Authors:** Yutaka Takaoka, Aki Sugano, Yoshitomo Morinaga, Mika Ohta, Kenji Miura, Haruyuki Kataguchi, Minoru Kumaoka, Shigemi Kimura, Yoshimasa Maniwa

**Affiliations:** Data Science Center for Medicine and Hospital Management, Toyama University Hospital, Toyama, Toyama 930-0194, Japan; Department of Computational Drug Design and Mathematical Medicine, Graduate School of Medicine and Pharmaceutical Sciences, University of Toyama, Toyama, Toyama 930-0194, Japan; Department of Medical Systems, Kobe University Graduate School of Medicine, Kobe, Hyogo 650-0017, Japan; Life Science Institute, Kobe Tokiwa University, Kobe, Hyogo 653-0838, Japan; Center for Clinical Research, Toyama University Hospital, Toyama, Toyama 930-0194, Japan; Department of Microbiology, Toyama University Graduate School of Medicine and Pharmaceutical Sciences, University of Toyama, Toyama 930-0194, Japan

**Author notes:** **Corresponding author:** Professor Yutaka Takaoka, Toyama University Hospital, Toyama, Toyama 930-0194, Japan, & Kobe University Graduate School of Medicine, Kobe, Hyogo 650-0017, Japan.

**Keywords:** SARS-CoV-2, COVID-19, Infectivity, Spike protein, Binding affinity, Mathematical model

## Abstract

Variants of a coronavirus (SARS-CoV-2) have been spreading in a global pandemic. Improved understanding of the infectivity of future new variants is important so that effective countermeasures against them can be quickly undertaken. In our research reported here, we aimed to predict the infectivity of SARS-CoV-2 by using a mathematical model with molecular simulation analysis, and we used phylogenetic analysis to determine the evolutionary distance of the spike protein gene (S gene) of SARS-CoV-2. We subjected the six variants and the wild type of spike protein and human angiotensin-converting enzyme 2 (ACE2) to molecular docking simulation analyses to understand the binding affinity of spike protein and ACE2. We then utilized regression analysis of the correlation coefficient of the mathematical model and the infectivity of SARS-CoV-2 to predict infectivity. The evolutionary distance of the S gene correlated with the infectivity of SARS-CoV-2 variants. The coefficient of the mathematical model obtained with results of molecular docking simulation also correlated with the infectivity of SARS-CoV-2 variants. These results suggest that the data from the docking simulation for the receptor binding domain of variant spike proteins and human ACE2 were valuable for prediction of SARS-CoV-2 infectivity. In addition, we developed a mathematical model for prediction of SARS-CoV-2 variant infectivity by using binding affinity obtained via molecular docking and the evolutionary distance of the S gene.

## 1. Introduction

Variants of the novel coronavirus (SARS-CoV-2 [severe acute respiratory syndrome coronavirus 2]) that are responsible for the worldwide pandemic known as COVID-19 have led to difficulties in enacting countermeasures against infection, because many variants have occurred in succession [1]. Infection control methods for SARS-CoV-2 have mainly included wearing face masks, avoiding close contact with other people (such as via lockdowns in urban areas), and providing multiple injections of vaccines [2]. The infectivity of SARS-CoV-2 variants has been rapidly changing, so understanding the infectivity of new variants is important for effective responses to the pandemic [3]. These changes have resulted in higher infectivities of new SARS-CoV-2 variants compared with past variants [4-9]. These alterations in infectivity indicate that the infectivity of new coronavirus variants may be estimated by utilizing the evolutionary distance of the spike protein gene (S gene) between the wild type and the variants. We previously described a mathematical model in which we used docking simulation results to predict UDP-glucuronosyltransferase 1A1 conjugation capacity [10]. By using the molecular docking simulation analyses, we found a plant leaf extract that inhibited the binding of SARS-CoV-2 spike protein to angiotensin-converting enzyme 2 (ACE2) [11]. Similarly, *in silico* docking data for a variant of SARS-CoV-2 spike protein and ACE2 may be used to estimate the infectivity of the new variant. In this research here, we choose these two approaches with mathematical models to achieve rapid and better understanding of the infectivity of new SARS-CoV-2 variants.

## 2. Materials and methods

### 2.1. Determination of the evolutionary distance between wild-type and variant S genes and infectivities of the variants

We chose six variants of SARS-CoV-2—alpha [12], beta [13], gamma [14], delta [15], omicron BA.1 [16], and omicron BA.2 [17]—for this research. We used the multiple sequence alignment method for the S gene of the SARS-CoV-2 variants, whose nucleotide sequences were obtained from NCBI, to perform an analysis via the ClustalW program [18]. FastTree [19] with default parameters then provided the phylogenetic tree and evolutionary distances between each mutant and wild type. The infectivity of each SARS-CoV-2 variant was obtained from previous research, and we summarized these data and developed an infectivity index for our research [4-9].

### 2.2. Analysis of the three-dimensional structures of SARS-CoV-2 spike protein, analysis of docking of the receptor-binding domain (RBD) and ACE2 protein, and development of a mathematical model

The amino acid sequence of wild-type SARS-CoV-2 spike protein was obtained from UniProt (UniProt ID: P0DTC2). The three-dimensional structure of SARS-CoV-2 spike protein (Protein Data Bank [PDB] ID: 6ZGG), which is an open state trimer, was used as the template for homology modeling of wild-type and variant SARS-CoV-2 spike proteins. We used Molecular Operating Environment software (Chemical Computing Group, Montreal, Quebec, Canada) to perform the homology modeling. The model structures were then subjected to structural optimization with molecular dynamics (MD) simulation, as described in our previous report [20]. After 10,000-ps MD simulation of each spike protein, docking analysis with ACE2 was performed as follows: The ACE2 structure was obtained from the PDB (PDB ID: 6M0J). To reduce the search space for docking, a partial structure of the RBD in the “up” conformation was cut from our simulated structure of the SARS-CoV-2 spike protein trimer. Docking sites were defined on the basis of the crystal structure of SARS-CoV-2 spike protein RBD bound with ACE2 (PDB ID: 6M0J). Two thousand docking runs were performed by using ZDOCK software (University of Massachusetts Medical School, Worcester, MA). To determine the stable complex group among the docking runs, the ZDOCK scores of the resulting complexes were clustered by using the group average clustering algorithm. The upper tail rule [21] was applied to determine the number of clusters. We used the most stable ZDOCK score in the docking runs and the average ZDOCK score of the most stable complex group to derive a mathematical model according to our previous research [10]. Figure 1 provides a flow diagram of this research.

**Figure 1.**
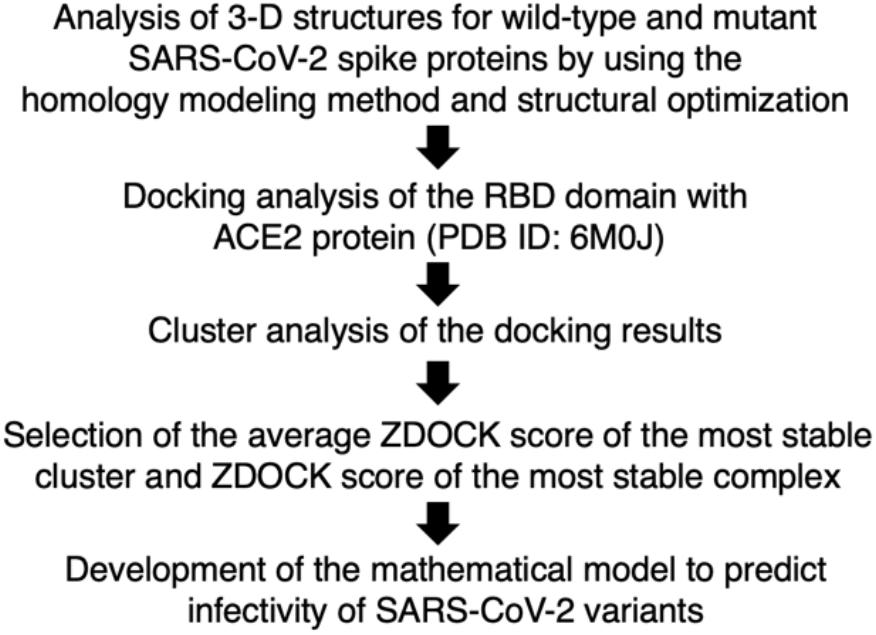
The method used for molecular simulation analysis.

## 3. Results

### 3.1. Evolutionary distances for S gene variants and results of docking of RBD with ACE2 protein

Table 1 shows the evolutionary distances between the wild type and the variants, as well as the docking affinities of RBD with ACE2. After multiple alignments of wild-type and variant S genes by using the ClustalW program, FastTree was used to determine the phylogenetic tree (data not shown) and evolutionary distances. The binding affinity of the SARS-CoV-2 spike protein RBD and ACE2 was determined by using the ZDOCK score as an indicator. ZDOCK scores were utilized in cluster analysis to determine the average ZDOCK score of the most stable cluster and ZDOCK score of the most stable complex for each variant. We used these results to develop a mathematical model to predict infectivities of SARS-CoV-2 variants.

**Table 1.**
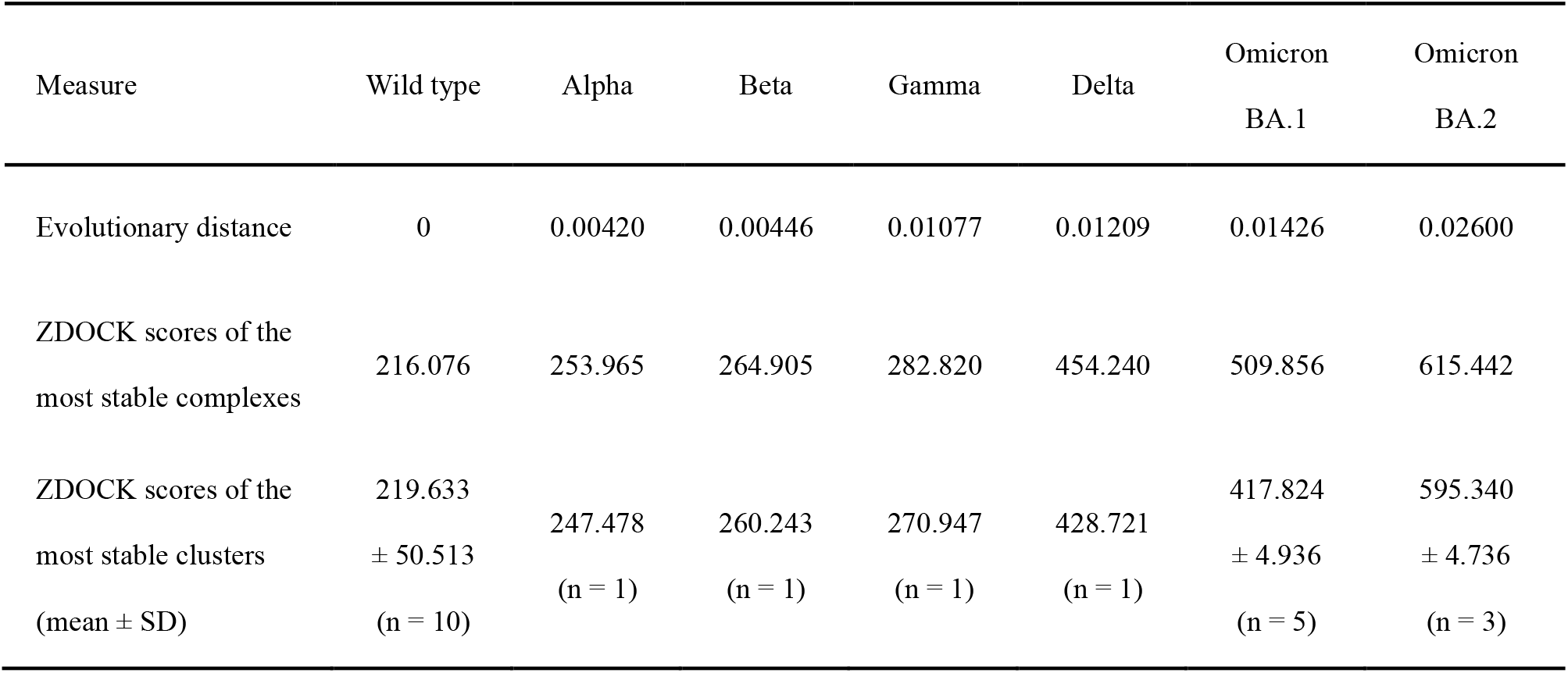
Evolutionary distances and binding affinities of SARS-CoV-2 spike proteins with ACE2.

### 3.2. Development of a mathematical model to estimate infectivities of SARS-CoV-2 wild type and variants

We developed a mathematical model to predict the infectivities of SARS-CoV-2 wild type and variants according to our previous research [10]. According to Fig. 2, the relationship between the evolutionary distance and infectivity of each variant can be represented by the following equation:

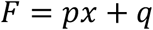

**Figure 2.**
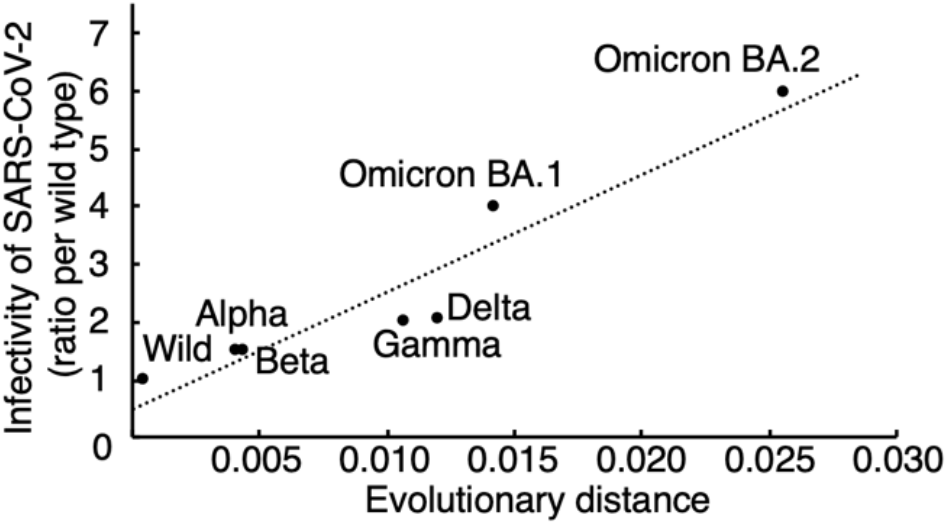
Correlation between evolutionary distance and infectivity of SARS-CoV-2 variants. Collinearity between the evolutionary distance and infectivity was noted. The regression line was indicated by dotted line as an equation of “y=201.06x + 0.5784.”

where *x* is the evolutionary distance between the wild-type (wild) and a mutant SARS-CoV-2 spike protein. Constant values *p* and *q* were estimated by minimizing the sum of squared error between the calculated infectivity and the reported infectivity (ratio per wild type) [4-9]. We then derived the mathematical model for the binding affinity of RBD and ACE2 by using the results of docking analyses. The coefficient of the mathematical model was derived from the following equation:

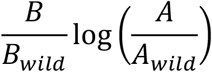

where A is the average ZDOCK score of the most stable cluster and B is the ZDOCK score of the most stable complex. The relationship between the coefficient of the variant infectivity and the reported infectivity (Fig. 3) can be represented by the following equation:

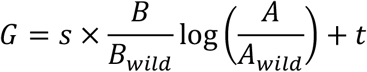

**Figure 3.**
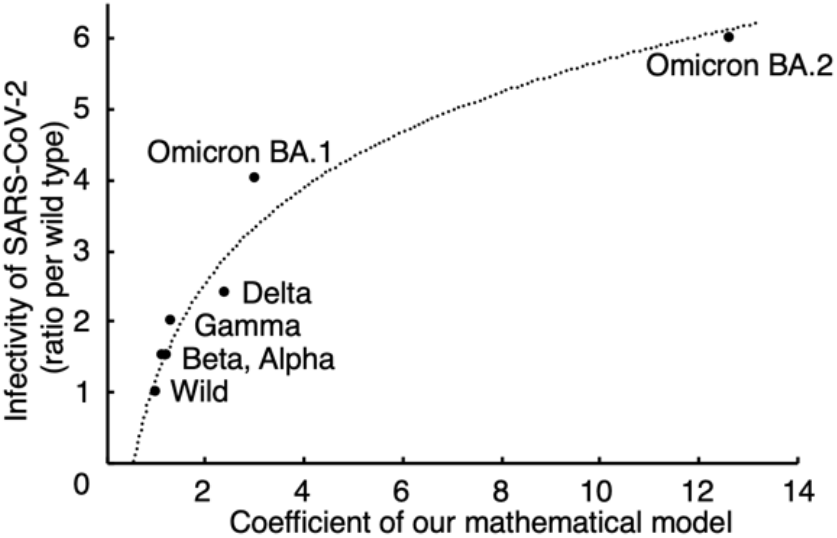
Correlation between infectivity of SARS-CoV-2 variants and the mathematical model based on molecular simulation analyses. The coefficient from our mathematical model with the molecular simulation data can be used to predict the infectivity of SARS-CoV-2 variants. The regression curve was indicated by dotted line as an equation of “y=1.9455ln(x) + 1.2029.”

Constant values *s* and *t* were estimated by minimizing the sum of squared error between the calculated infectivity and the reported infectivity (ratio per wild type) [4-9]. The infectivity of SARS-CoV-2 (*P*) is defined by the evolutionary distance and the docking result with ACE2:

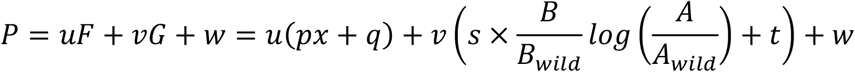

Constant values *u, v*, and *w* were estimated by minimizing the sum of squared error between the calculated infectivity P and the reported infectivity of each SARS-CoV-2 variant *V*:

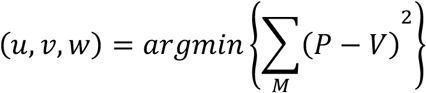

where set *M* represents the SARS-CoV-2 variants whose infectivities are reported. A correlation was seen between the coefficient of the mathematical model and the evolutionary distance of SARS-CoV-2 spike protein (data not shown). These results suggest that our mathematical model can predict the infectivities of SARS-CoV-2 wild type and variants. Finally, the infectivity of SARS-CoV-2 variants is defined by the following equation:

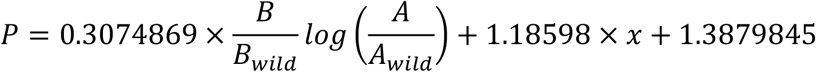

## 4. Discussion

Reported infectivities of SARS-CoV-2 variants were correlated with both evolutionary distance of the S gene and the coefficient of the mathematical model according to our *in silico* docking data (Figs. 2 and 3). With our mathematical model, these two factors reproduced the reported infectivity better than the model using only one factor in the coefficient of determination (data not shown).

In previous research, only the RBD domain of the spike protein was used in the analysis of binding affinity with ACE2: no MD analysis of the structure optimization [22-26] and/or local optimization of only the RBD domain [22-28] of the spike protein. However, we chose the three-dimensional structure of the complete trimeric spike protein (PDB ID: 6ZGG) in homology modeling in this research. After MD analysis, we confirmed the trajectory of the root mean square deviation and the disallowed regions of the Ramachandran plot for proper quality of each variant spike protein structure. The RBD domain of the variant trimeric spike protein structures was then provided for the docking analyses with ACE2. Therefore, our experimental strategy achieved high accuracy because we used a molecular simulation method which possibly imitates the behavior of the biomolecules. In addition, our previous research [29-34] suggested the value of this prediction that utilized our molecular simulation methods. Additional analyses of binding affinities of other spike proteins of SARS-CoV-2 variants with ACE2 may provide greater accuracy, and such experiments are now in progress.

## Abbreviations

S gene: spike protein gene
ACE2: angiotensin-converting enzyme 2
RBD: receptor-binding domain
MD: molecular dynamics
PDB: Protein Data Bank

## Data availability

Data that support the findings of this study are available from the corresponding author upon reasonable request.

## Author contributions

Y.T. conceived and designed this research. Y.T. and S.A. preformed the analyses and acquired the data. Y.T., S.A., Y.M., and M.O. interpreted the data. Y.T., S.A., and K.M derived mathematical model. Y.T. and S.A. wrote the draft, and all authors reviewed and approved the manuscript.

## Acknowledgements

This work was supported by JSPS Grant-in-Aid for Scientific Research grant numbers 21K12110 to Y.T. and 19K12202 to A.S.

